# Single-cell transcriptomics of the mouse gonadal soma reveals the establishment of sexual dimorphism in distinct cell lineages

**DOI:** 10.1101/410407

**Authors:** Isabelle Stévant, Françoise Kühne, Andy Greenfield, Marie-Christine Chaboissier, Emmanouil T. Dermitzakis, Serge Nef

## Abstract

Sex determination is a unique process that allows the study of multipotent progenitors and their acquisition of sex-specific fates during differentiation of the gonad into a testis or an ovary. Using time-series single-cell RNA sequencing (scRNA-seq) on ovarian Nr5a1-GFP^+^ somatic cells during sex determination, we identified a single population of early progenitors giving rise to both pre-granulosa cells and potential steroidogenic precursor cells. By comparing time-series scRNA-seq of XX and XY somatic cells, we demonstrate that the supporting cells emerge from the early progenitors with a non-sex-specific transcriptomic program, before pre-granulosa and Sertoli cells acquire their sex-specific identity. In XX and XY steroidogenic precursors similar transcriptomic profiles underlie the acquisition of cell fate, but with a delay in XX cells. Our data provide a novel framework, at single-cell resolution, for further interrogation of the molecular and cellular basis of mammalian sex determination.

## Introduction

Testes and ovaries have the same developmental origin: the gonadal primordia. These start developing around embryonic day (E)9.5 in mice through the thickening and proliferating coelomic epithelium on the ventromedial surface of the mesonephroi (Byskov, 1986). Before sex determination, the gonadal primordia, also called bipotential gonads, are composed of multipotent somatic progenitor cells that are competent to adopt one or the other sex-specific cell fate, and of migrating primordial germ cells. Sex determination is initiated in the supporting cell lineage around E11.5, leading to their differentiation as Sertoli cells in XY gonads following the expression of *Sry* (Albrecht and Eicher, 2001; Sekido et al., 2004) or as pre-granulosa cells in XX with the stabilisation of WNT/β-catenin signalling (Chassot et al., 2008, 2012). Following supporting cell differentiation, this sex fate decision propagates to the germ cells and the other somatic lineages, including the steroidogenic cells (Leydig cells in XY, and theca cells in XX) that later drive the primary and the secondary sexual characteristics through hormonal control.

While the origin of germ cells is well established (for review, see (De Felici, 2016)), the origins of somatic cell lineages of both the XX and XY gonads is not as well understood. With respect to supporting cells, lineage tracing experiments indicate that Sertoli cells and pre-granulosa cells arise from common supporting precursor cells competent to express *Sry*, and originating from the coelomic epithelium (Albrecht and Eicher, 2001; Karl and Capel, 1998; Mork et al., 2012). Unlike the Sertoli cell population that expands by mitotic division until puberty, the granulosa cell population expansion is more complex and not fully resolved. Pre-granulosa cells are mitotically inactive and expand in two phases by the differentiation of ingressing coelomic epithelial cells (Gustin et al., 2016). The first wave of granulosa cell recruitment occurs in foetal life between E11.5 and E14.5 and contributes to the medullary follicles that are activated at birth immediately after their assembly (Mork et al., 2012). The second wave of granulosa cell recruitment occurs at birth and contributes to the cortical primordial follicles. By post-natal day (P)7, the follicle assembly is almost complete and no new granulosa cells are recruited (Mork et al., 2012).

In respect of the steroidogenic cell lineage, the foetal Leydig cells arise from precursor cells present in the interstitial compartment of the testis originating from N5RA1-expressing cells derived from the coelomic epithelium (Karl and Capel, 1998; Stévant et al., 2018) and also from migrating mesonephric cells (DeFalco et al., 2011; Rotgers et al., 2018). However, the origin of theca cells, the female counterpart to Leydig cells, is still unclear. Theca cells arise in the mouse ovary during the first ten postnatal days by the recruitment of uncharacterised precursor cells through Hedgehog signals secreted by the granulosa cells while follicles enter their secondary stage (Liu et al., 2015; Young and McNeilly, 2010). One suspected source of theca cell precursors resides in a stromal cell population present in the foetal ovary that expresses MAF (or c-MAF) and MAFB (DeFalco et al., 2011; Jameson et al., 2012), and NR2F2 (also known as COUP-TFII) (Rastetter et al., 2014; Takamoto et al., 2005). Moreover, a recent lineage tracing study showed that steroidogenic theca cells mainly derive from cells expressing WT1 at E10.5, and to a lesser extent from mesonephric cells expressing GLI1 at E12.5 (Liu et al., 2015). Beyond the origin of the supporting and the steroidogenic cell lineages in both male and female gonads, our knowledge of the mechanisms of the specification of the cell lineages into their respective sex-specific fate remains incomplete.

In order to improve our understanding of somatic cell lineage specification and sex-specific cell differentiation during the process of sex determination, we performed time-series single-cell RNA sequencing (scRNA-seq) of *Nr5a1*-GFP somatic cells composing the developing XX gonad from E10.5 to as late as P6, and we reconstructed the cell lineage specification through time, as previously described with the fetal testis (Stévant et al., 2018). By combining the expression data from both XX and XY time-series scRNA-Seq, we compared the transcriptomic profiles of the supporting and the interstitial/stromal *Nr5a1*-GFP somatic cells of the two sexes as they differentiate and demonstrated that the supporting cells differentiate in two steps, while the interstitial/stromal cells display a very similar specification toward a steroidogenic fate.

## Results

### XX somatic cells are classified in four transcriptionally distinct cell populations

To characterise and reconstruct the somatic cell lineages of the developing XX gonad as ovary development proceeds, we purified the somatic cells at six different developmental stages of gonadal differentiation (E10.5, E11.5, E12.5, E13.5, E16.5 and P6) using the *Tg(Nr5a1-GFP)* transgenic mouse (**Figure 1A**) (Stallings et al., 2002). *Nr5a1* is expressed specifically in gonadal somatic cells giving rise to the supporting and the steroidogenic lineages, in both male (XY) and female (XX) gonads, and in a time window large enough to cover the bipotential state prior to sex determination and the whole of gonadal development (Nef et al., 2005; Stévant et al., 2018). Thus, the *Tg(Nr5a1-GFP)* transgenic mouse constitutes a powerful tool for isolating gonadal somatic cells and studying their differentiation (**Figures 1B-C** and **Figure S1**). Briefly, at each relevant embryonic stage, gonads from *Tg(Nr5a1-GFP)* animals were dissociated and the *Nr5a1*-GFP^+^ cells were sorted by fluorescent active cell sorting (FACS) (**Figure S1**). GFP^+^ cells were isolated and processed with the Fluidigm C1 Autoprep system and sequenced (full-length RNA-seq, 100bp paired-end reads, 10M reads per cell) (**Figure 1D**). A total of 563 cells remained after filtering based on various quality control metrics (see **Methods**).

**Figure 1:**
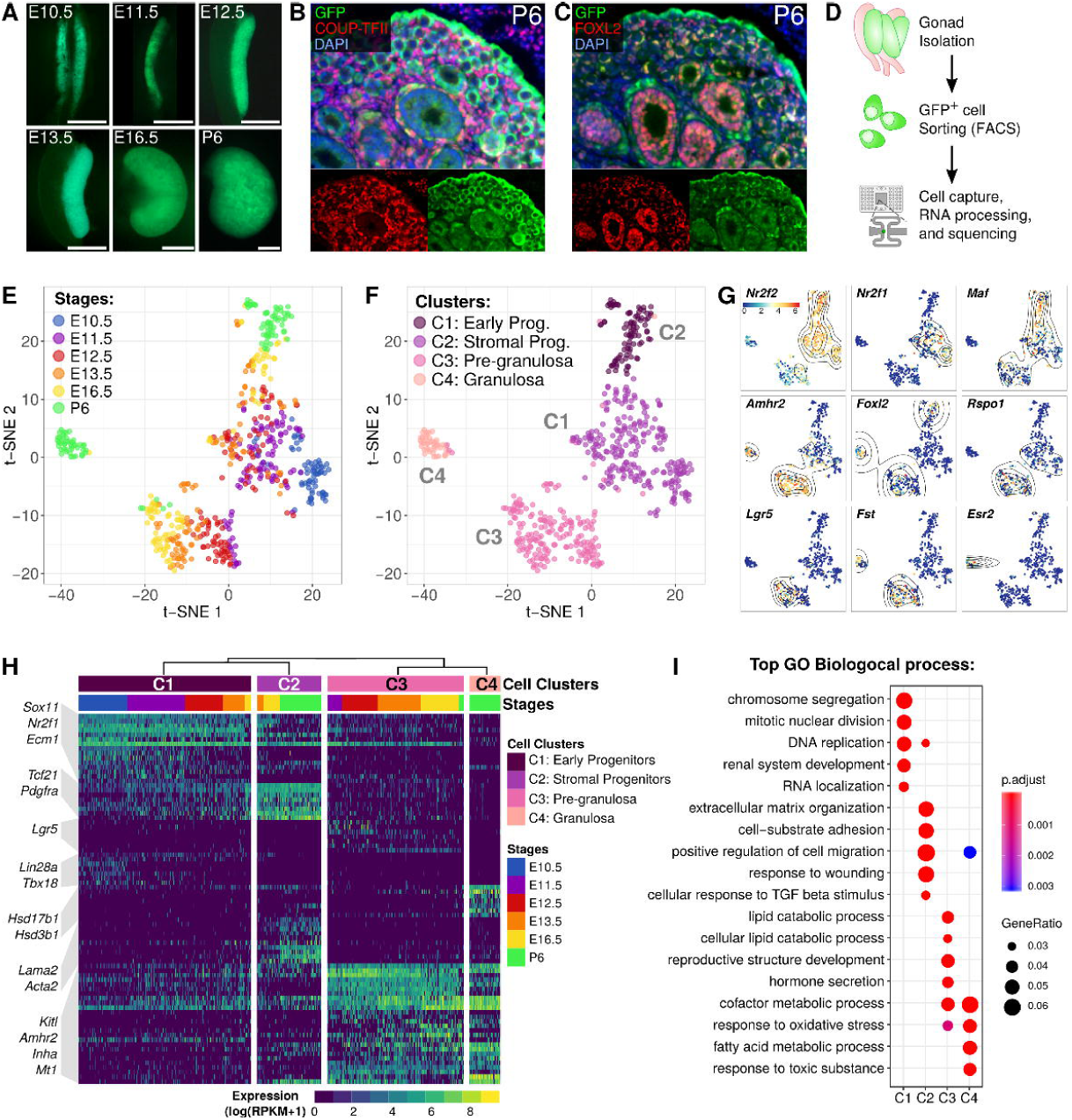
Classification and identification of the XX gonadal somatic cells during early ovarian development. (A) Images of XX *Nr5a1*-GFP^+^ gonads at six stages of development under UV light (scale bars=500μm). (B) and (C) Co-immunofluorescence of GFP and marker genes for stromal progenitors (NR2F2), and granulosa cells (FOXL2) at P6. (D) Experimental design. *Nr5a1*-GFP^+^ cells from XX gonads were sorted by FACS prior to the single-cell capture, and RNA sequenced. (E) and (F) Two-dimensional t-SNE (t-distributed stochastic neighbour embedding) representation of the 563 XX GFP^+^ single-cells. Cells are coloured by embryonic stages (D) and by cell clusters (E). (G) T-SNE representations with cells coloured by the expression level of marker genes of the ovary. The black lines represent the spatial density of the cells expressing the given gene higher than the mean level of expression. (H) Heatmap showing the expression of the top 20 differentially expressed genes for each cell cluster (q-val<0.05). (I) Top results from Gene Ontology (GO) enrichment test showing the terms associated with the up-regulated genes from each of the four cell clusters.

We classified the different somatic cell populations present in the developing ovary using the same method we developed for the analysis of testis development, *i.e.* we selected the highly variable genes, performed a principal component analysis (PCA) and hierarchical clustering on the significant principal components (see (Stévant et al., 2018) and **Methods**). We obtained four cell clusters combining different embryonic stages (clusters C1 to C4) (**Figures 1E &F**). Expression enrichment of known markers and differentially expressed genes (**Figures 1G &H**, **Figure S2**, **Supplementary Data 1**), allowed us to assign the identity of the cell clusters.

The clusters C1 and C2 represent *Nr2f2*-expressing progenitor cells. Cluster C1 forms the early progenitor cell population, with cells from E10.5 to E13.5, and expresses markers found in our previous study such as *Nr2f1* (Stévant et al., 2018); cluster C2 contains cells from E13.5 onward, which are stromal progenitor cells that express the stromal marker gene *Maf* (DeFalco et al., 2011) (**Figure 1G**), and surprisingly some markers of foetal Leydig cell progenitors, *Tcf21* and *Pdgfra* (Brennan et al., 2003; Cui et al., 2004; Stévant et al., 2018) (**Figure 1H**).

In contrast, cells from clusters C3 and C4 co-express the supporting cell marker *Amhr2* (Baarends et al., 1995), *Foxl2* (Mork et al., 2012; Schmidt et al., 2004; Uda et al., 2004), and *Fst* (Bouma et al., 2007), and represent granulosa cells at different stages of their differentiation (**Figures 1G &H**). Cluster C3 contains foetal cells from E11.5 to E16.5 and some P6-expressing genes related to pre-granulosa cells, such as *Lgr5* (**Figure 1H** & **Figure S2**). The C4 cluster contains mostly post-natal P6 cells that express oestrogen receptor *Ers2*, and the steroid-related genes *Hsd3b1* and *Hsd17b1*. Therefore, we named the C3 cluster ‘pre-granulosa’, and the C4 cluster ‘granulosa’, to distinguish these two populations.

Over-represented GO terms associated with the differentially expressed genes in each cell cluster (**Figure 1I**, **Supplementary data 2**) reveal that the early progenitor cells are proliferating cells, whereas the stromal progenitors express genes related to morphological organisation (“extracellular matrix organisation”, “positive regulation of cell migration”). The pre-granulosa and granulosa cells express genes related to lipid metabolic processes and “hormone secretion”, indicating that these cells have the capacity to produce hormones as early as foetal life (Dutta et al., 2014).

To summarise, the time-series single-cell RNA-seq of the *Nr5a1*-GFP^+^ cells of the developing XX gonad from E10.5 to P6 identified four cell populations, including early and stromal progenitor cells, pre-granulosa cells and post-natal granulosa cells. The absence of other detected cell types at P6 indicates that we did not capture theca cells; these might constitute a rare cell population at that developmental stage.

### Cell lineage reconstruction identifies the dynamics of gene expression during ovarian fate commitment

The reconstruction of the *Nr5a1*-GFP^+^ cell lineages in the developing XX gonad allowed us to identify transition states leading to the differentiation of the granulosa cells and the stromal cells from a common progenitor cell population (**Figures 2A-B**, **Methods**). We observe that the early progenitor cells give rise subsequently to the granulosa cell lineage and the stromal progenitor cell lineage around E11.5-E12.5.

**Figure 2:**
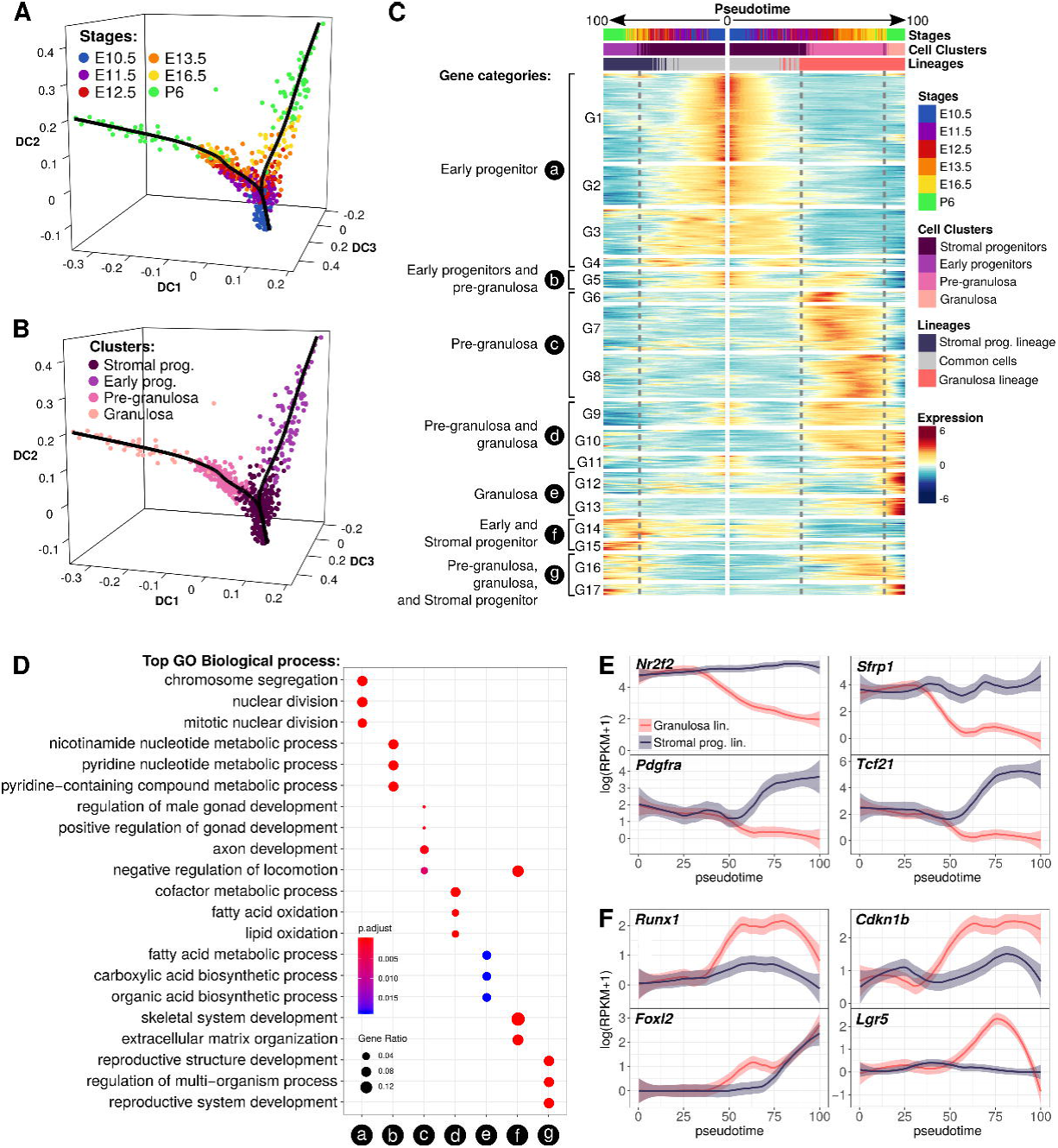
Cell lineage reconstruction and identification of the genetic program driving granulosa and stromal cells. (A) and (B) Diffusion map of the most variable genes and reconstruction of the cell lineages. Dots represent cells and the black lines represent the estimated cell lineages. (A) is coloured by embryonic stages and (B) by cell clusters. (C) Heatmap representing the dynamics of gene expression in the progenitor (left) and granulosa cell (right) lineage through the pseudotime. Early progenitor cells common to both lineages are located in the centre (0) of the map with the divergence of granulosa cells to the right and evolution of progenitor cells to the left. Gene expression was normalised to the mean, and classified with k-means (k=17). They were categorised according to the cell types in which they are overexpressed (a-g). The dotted lines mark the limit of the cell clusters. (D) Top results from Gene Ontology (GO) enrichment test showing the terms associated with the gene categories defined in (C). (E) and (F) Expression profiles of genes that become restricted to stromal progenitors (E) or granulosa cells (F). The solid line represents the loess regression, and the fade band is the 95% confidence interval of the model.

By ordering the cells along a pseudotime (Street et al., 2018), we identified genes that are dynamically expressed during the specification of the two cell lineages (differentially expressed genes along the pseudotime, *q-value*<;0.05) (**Figure 2C**, **Methods**). A total of 3,916 genes reveal a dynamic expression profile through time, with 1,733 restricted to the granulosa cell lineage, 1,059 to the stromal progenitor cell lineage, and 1,124 dynamically expressed in the two cell lineages (**Supplementary Data 3**). We represented the smoothed gene expression level (loess regression) of the two cell lineages with a double heatmap, where the centre represents the starting point of the lineage (Pseudotime 0), and the extremities represent the lineage end-points (Pseudotime 100) of the stromal progenitors (left) and the granulosa cell lineages (right), respectively (**Figure 2C**). We classified these genes by expression patterns (G1-G17), and by cell-specific expression categories (a-g) to eventually identify enrichment of biological processes through a GO term over-representation test (**Figure 2D, Supplementary Data 4**). The heatmap revealed that granulosa cells diverge from the early progenitor cells with a strong differentiation program compared to the stromal progenitor cells, which exhibit many fewer dynamic genes.

The genes that are over-expressed in the common progenitor cells of the undifferentiated gonad (patterns G1 to G4, or category (a), **Figure 2C**) are related to mitotic cell division, mesonephros development, positive regulation of stem cell development and epithelium morphogenesis, consistent with their coelomic epithelial origin (**Figure 2D**, **Supplementary Data 4**). Their expression level decreases during commitment to both pre-granulosa and stromal progenitor fate, suggesting a cell identity conversion. We noted that this cell conversion occurs around E11.5-E12.5 in the granulosa lineage, while it occurs from E13.5 in the stromal progenitor lineage (**Figures 2A-C**). This suggests that the progenitor cells remain competent to be recruited as pre-granulosa cells as late as E12.5.

The differentiation of the granulosa cell lineage is characterised by highly dynamic transcriptomic profiles involving transient and permanent activation of genes. Genes over-expressed in pre-granulosa cells only (category (c), 608 genes) are classified in three expression profiles (G6 to G8) (**Figure 2C** and **Supplementary data 3**). Of interest, the G6 profile is composed of 62 genes that are transiently over-expressed at E11.5 and E12.5, at the onset of pre-granulosa cell differentiation. Among these genes, we found the male testis-maintenance gene *Dmrt1* (Lei et al., 2007), *Cyp11a1*, as previously observed in the pre-Sertoli cells (Stévant et al., 2018; Val et al., 2007), and *Lgr4*, known to act as a receptor of RSPO1 and a promoter of Wnt/beta-catenin signalling (Koizumi et al., 2015) (**Supplementary Data 3**). The profiles G9 to G11, or category (d), contain 368 genes that are expressed as soon as the pre-granulosa cells differentiate and are maintained after birth (**Figure 2C**). These genes are mostly related to lipid metabolic processes (**Figure 2D** and **Supplementary Data 4**). In this category we also found genes known as involved in ovarian development, such as *Kitl* (Hutt et al., 2006; Jones and Pepling, 2013), *Fst* (Kashimada et al., 2011), and the steroidogenic acute regulatory protein *Star* (Caron et al., 1997). And finally, the profiles G12 and G13, or category (e), contain 242 genes that are over-expressed in the post-natal granulosa cells at P6 (**Figure 2C** and **Supplementary Data 3**). This includes genes which has been previously described as expressed in foetal Sertoli cells such as *Amh*, which is also known to control primordial follicle recruitment (Durlinger et al., 1999, 2002), *Aard* (Bouma et al., 2010), and *Mro* (Smith et al., 2003, 2008). We also found the expression of *Igf1r* (Baumgarten et al., 2017), which is required for steroidogenesis, and *Inha* and *Inhbb* (Findlay, 1993; Mather et al., 1997; Weng et al., 2006) (**Supplementary Data 3**).

In contrast to the highly dynamic program mediating granulosa cell differentiation, the progenitor cell lineage displays much less variation in gene expression during the process of ovarian development. The gene category (f) regroups 173 genes that are expressed in the stromal progenitor lineage, with the pattern G14 containing genes expressed as early as E10.5 and over-expressed at P6, and the pattern G15, which contains genes over-expressed at P6. Among them, we found genes known as markers of steroidogenic cell precursors such as *Wnt5a* (Stévant et al., 2018), *Pdgfra* (Brennan et al., 2003), *Tcf21* (Bhandari et al., 2012; Cui et al., 2004), *Gli2* (Barsoum and Yao, 2011), *Arx* (Miyabayashi et al., 2013), and the secreted negative regulator of the *Wnt* signalling pathway, *Sfrp1* (Warr et al., 2009). These results suggest that the progenitor cell lineage undergoes transcriptional changes that restrict its competence towards a steroidogenic fate required for the differentiation of theca cells.

Overall, the reconstruction of the *Nr5a1*-GFP^+^ cell lineage reveals that before female sex determination, there exists a single, highly proliferative progenitor cell population. A subset of this population differentiates, firstly, as pre-granulosa cells and, ultimately, as granulosa cells, driven by a dynamic transcriptional program composed of the transient expression of hundreds of genes, including genes implicated in the *Wnt* signalling pathway. Conversely, the remaining progenitor cell population exhibits transcriptomic changes that restrict its competence towards a steroidogenic fate during foetal ovarian development, ultimately expressing genes known as makers of steroidogenic cell precursors in the testis.

### Integrating scRNA-seq of both XX and XY Nr5a1-GFP^+^ cells reveals the establishment of sexual dimorphism

By combining scRNA-seq data from 400 XY *Nr5a1*-GFP^+^ cells (Stévant et al., 2018) with the present data from 563 XX *Nr5a1*-GFP^+^ cells, we undertook to investigate the transcriptomic programs of sex determination and characterize the sex-specific differences in each cell lineage.

We merged the two datasets and applied the same clustering method as previously, and we obtained five major cell clusters (clusters D1 to D5) (**Figures 3A-C**, **Methods**). The early progenitor populations from XX and XY gonads cluster together (cluster D1), as well as the XX stromal and the XY interstitial progenitors (Cluster D2), even though we observe a tendency to segregate by sex in the t-SNE representation (**Figures 3C**). This suggests that the progenitor cell lineages of both XX and XY gonads do not display sufficient sexual dimorphism to permit segregation and be considered as different cell types, even late in development. Interestingly, the cluster D3 contains pre-granulosa cells from E11.5 to P6 but also E11.5 pre-Sertoli cells. The clusters D4 and D5 contain granulosa cells at P6, and Sertoli cells from E12.5 to E16.5, respectively. This suggests that the supporting cells emerge from the progenitors with a similar transcriptomic program, and that Sertoli cells differentiate with more pronounced and dynamic transcriptomic changes when compared to pre-granulosa cells, which complete their differentiation after birth.

**Figure 3:**
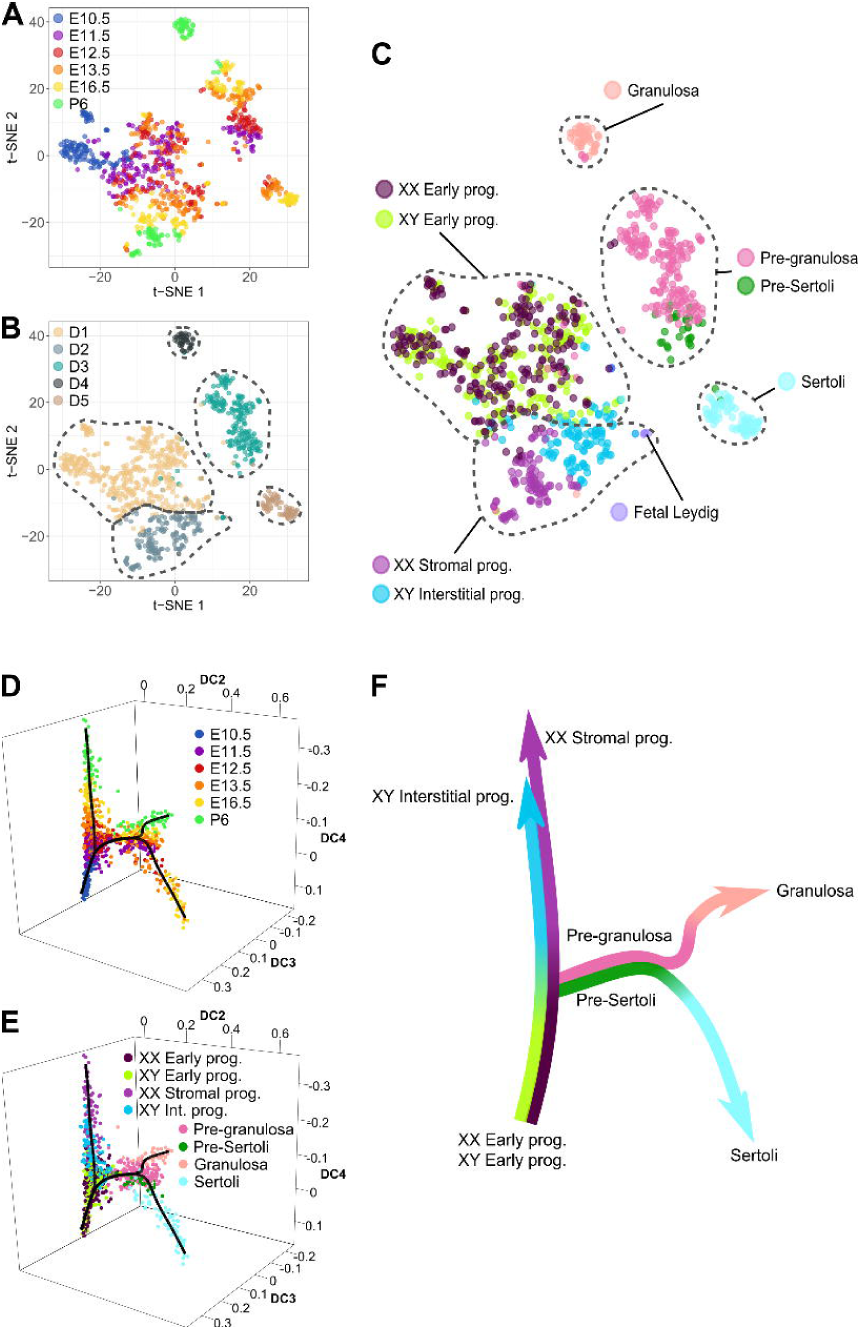
Progression of the gonadal somatic cell transcriptomes in both sexes during sex determination. (A), (B) and (C) T-SNE representation of the 963 single cells from XX and XY developing gonads coloured by the cell clustering coloured by embryonic stages (A), by cell clusters (B), and by cell types identified in the sex-respective analysis (C). (D) and (E) Diffusion map and reconstruction of the cell lineages coloured by embryonic stages (D), and by cell types (E). (F) Simplified representation of the cell lineage reconstruction.

The diffusion map and the lineage reconstruction (**Figures 3D-F**) confirms what was observed using clustering. The progenitor cell lineage combines both sexes from E10.5 to late stages (**Figures 3D-E**), and the supporting cell lineage diverges from progenitors independently of genetic sex, and subsequently gives rise to Sertoli and granulosa cells with distinct timings (E12.5 in XY, and E16.5-P6 in XX).

### Commitment to the supporting cell lineage involves a common intermediate differentiation step prior to the emergence of sexual dimorphism

We analysed how the supporting cell lineage emerges and acquires its respective sex-specific cell types. We first evaluated when the supporting cell lineage transcriptomes become sexually dimorphic. To do so, we used cell lineage reconstruction to divide the supporting cell lineage into three developmental windows defined by the branch points (BP) of the lineage trajectories (**Figure 4A**) and we determined whether genes previously identified as dynamically expressed (from **Figure C2** and from (Stévant et al., 2018)) display sexual dimorphisms within these windows (differentially expressed genes, q-values<;0.05, **Supplementary data 5**). We conclude that sexual dimorphism appears in the supporting cell lineage after branch point 2, which corresponds to the transition between pre-Sertoli and Sertoli cells from E12.5 in XY, and to pre-granulosa cells from E13.5 in XX (**Figure 4A & B**).

**Figure 4:**
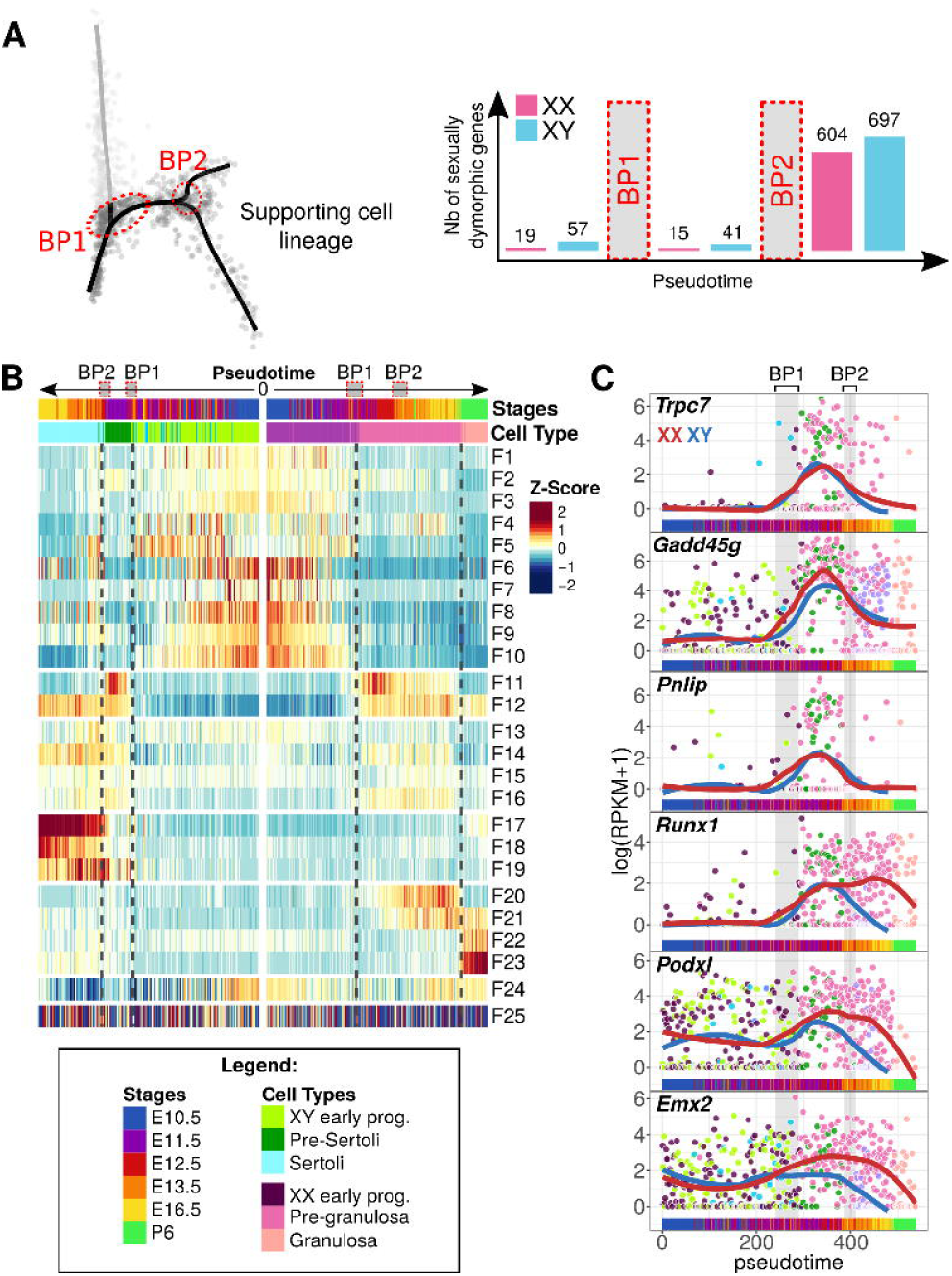
Emergence of sexual dimorphism in the supporting cell lineage. (A) Sexual dimorphic genes during three phases of the supporting cell lineage specification, *i.e.* prior to (before branch point (BP) 1), during (between BP1 and BP2) and after (after BP2) supporting cell commitment. (B) Heatmap representing the transcriptomic progress of the Sertoli and the granulosa cell lineage dynamic genes through the pseudotime. Genes were grouped by expression profiles with k-means. The heatmap shows the normalized average z-score of each gene group. Early progenitor cells common to both lineages are located in the center (0) of the map with divergence of Sertoli cells to the left and granulosa cells to the right. The dotted lines mark the limit of the cell clusters. (C) Expression profiles of genes showing an over-expression at the onset of supporting cell commitment. The solid lines represent the loess regression of the expression for each sex.

We then compared the expression dynamics of Sertoli and granulosa cell differentiation by classifying genes according to expression profiles in both sexes (k-means, k=25) (**Supplementary data 5**), and we represented on a double heatmap the averaged z-scores of each of the expression profile clusters (F1-F25) (**Figure 4B**). We see that the genes expressed in early progenitor cells are down-regulated during the commitment of the supporting cells of both sexes (gene profiles F1 to F10, **Figure 4B**). Then, we observed transient over-expression of 168 genes at the onset of both pre-Sertoli and pre-granulosa cell differentiation (gene profiles F11, **Figure 4B & C**). These genes represent a narrow window of expression in E11.5 pre-Sertoli cells, while they are over-expressed from E11.5 to E16.5 in the pre-granulosa cells. Among them, only *Sry* is significantly sexually dimorphic between BP1 and BP2 (**Supplementary data 5**). In this expression profile, we also found *Gadd45g*, which controls *Sry* expression via MAPK signals (Gierl et al., 2012; Warr et al., 2012), *Runx1* (Munger et al., 2013; Nef et al., 2005), *Podxl* (Herrera et al., 2005) or the early gonadal ridge promotor genes *Emx2* and *Lhx9* (Birk et al., 2000; Miyamoto et al., 1997) (**Figure 4C** and **Supplementary data 5**). These data suggest that pre-granulosa cells remain in a progenitor-cell-like state during foetal life and complete their differentiation after birth.

With this analysis, we also observed a considerable number of genes that share the same expression behaviour during differentiation of Sertoli and granulosa cells (gene profiles F12 to F16, 931 genes, **Figure 4A**), and also genes that become sexually dimorphic (gene profiles F17 to F24, 1071 genes), from E12.5 in Sertoli cells, and from E13.5 in pre-granulosa cells.

These results suggest that supporting cells undergo commitment from the early progenitor cells at around E11.5 with a common genetic program that is not strictly related to sex determination, despite the expression of *Sry*. This is consistent with the fact that XX supporting cell precursors are competent to express *Sry*, as shown by the activation of an *Sry*-YFP transgene in XX gonads (Albrecht and Eicher, 2001; Harikae et al., 2013; Koopman et al., 1991). This precise moment constitutes the bipotential state of the supporting cell lineage, after having diverged from the progenitor cell lineage. We also noted that in XX gonads, the bipotential state of supporting cells lasts one day longer than in XY gonads, reflecting the rapid effect of *Sry* on the activation of the Sertoli cell differentiation program.

### Stromal and interstitial cells display modest sexual dimorphism but differ in timing of expression

We repeated our analysis of the progenitor cell lineage to examine the emergence of sexual dimorphism as the XY interstitial and the XX stromal cells progress during gonadal development. We observe that the progenitor cells display modest sexual dimorphism after the branch point 1, with 117 genes over-expressed in XY (**Figure 5A** and **Supplementary data 6**). When we examine the dynamics of gene expression in XX and XY progenitors side by side (**Figure 5B**), we see that the same genes are up-regulated in both sexes while cells progress from the early progenitor state to stromal and interstitial cells, respectively, including markers of XY steroidogenic cells, *Pdgfra, Arx* and *Ptch1* (**Figure 5C** and **Supplementary data 6**). However, we note that the timing of gene expression is not the same, such as *Ar* and *Acta2*, which are expressed from E13.5 in XY interstitial cells and from P6 in XX stromal cells (**Figure 5C**). We also found a few genes that are exclusive to one or the other sex, such as *Inhba* (H19), which is expressed in XY interstitial cells only, and *Foxl2* (H23), which is expressed in P6 XX stromal cells (**Figure 5B, C** and **supplementary data 6**).

**Figure 5:**
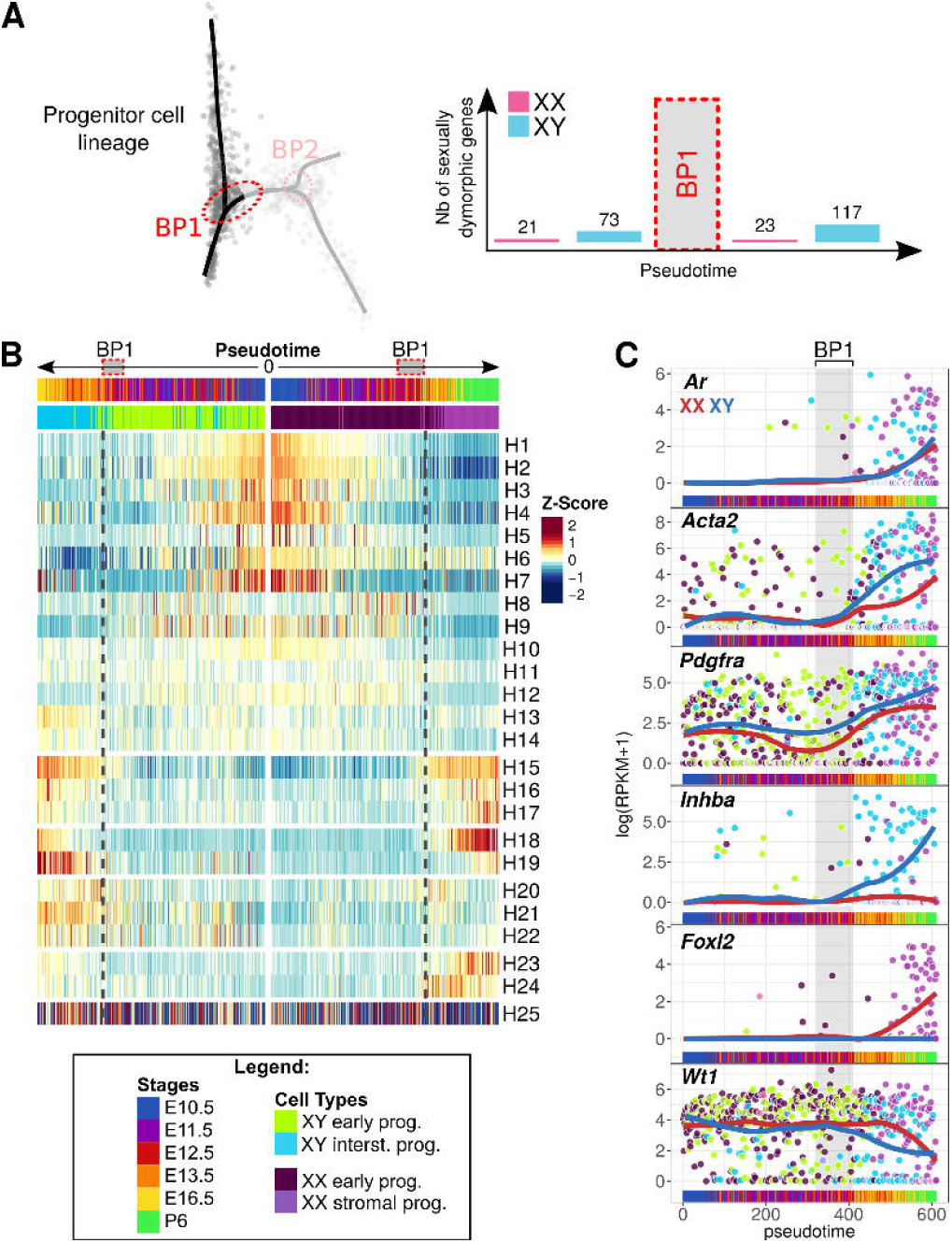
Progression of the interstitial/stromal progenitor cells. (A) Sexually dimorphic genes during the two phases of the progenitor cell lineage specification, *i.e.* prior to BP1, and after BP1. (B) Heatmap representing the transcriptomic progress of the interstitial and the stromal cell lineage dynamic genes through the pseudotime. Genes were grouped by expression profiles with k-means. The heatmap shows the normalized average z-score of each gene group. Early progenitor cells common to both lineages are located in the center (0) of the map with divergence of interstitial cells to the left and stromal cells to the right. The dotted lines mark the limit of the cell clusters. (C) Expression profiles of genes showing interesting profiles of expression. The solid lines represent the loess regression of the expression for each sex.

These results suggest that the progenitor cell lineage commits to the steroidogenic fate earlier in the XY gonad, around E12.5, to allow these cells to differentiate as foetal Leydig cells, whereas XX stromal cells commit to steroidogenic progenitor cells from E16.5, to ultimately differentiate as theca cells after birth, when granulosa cells also complete their differentiation.

## Discussion

With this study, we aimed to analyse the transcriptomic programs at play during sex determination in *Nr5a1*^+^ somatic cells of XX and XY mouse gonads. Subsequent to our previously published analysis of the early testis development (Stévant et al., 2018), we sequenced and analysed the transcriptome of hundreds of individual XX *Nr5a1*-GFP^+^ gonadal somatic cells, from the bipotential gonads at E10.5, to post-natal ovaries at P6. By combining these transcriptomic data from XX and XY cells, we were able to reconstruct the sequence of transcriptomic events underlying the cell lineage specification and the sex-specific cell differentiation of both the supporting cell lineage and the steroidogenic precursor cell lineage from a common multipotent progenitor cell population.

In contrast to the testis, foetal ovarian development is characterized by modest and late morphological changes; as a consequence, early ovarian development was not understood so well. The cell type contribution of the expression of critical factors such as *Fst, Wnt4* or *Rspo1*, remained unclear in the absence of cell lineage tracing. Our unsupervised analyses of XX *Nr5a1*-GFP^+^ somatic cells from E10.5 to P6 allow the identification of the most abundant somatic cell populations present during early ovarian development and the study of their lineage specification, their transcriptomic signatures, and the expression dynamics as they differentiate. We found a similar lineage specification pattern to that observed during early testis development (Stévant et al., 2018), with the presence of a common progenitor cell population at E10.5, from which differentiate the pre-granulosa cells. The remaining progenitor cells that do not commit as pre-granulosa express *Nr2f2* at high levels and persist in the ovary until as late as P6. We were able to identify the transcriptomic signature of the first commitment of pre-granulosa cells from early progenitors from E11.5 onward; instead, we showed that the transcriptome of pre-granulosa cells changes throughout foetal life, with successive activation and repression of hundreds of genes. The progenitor cell lineage that remains in the stroma displays progressive transcriptomic changes, at around E13.5, which specify their identity as potential steroidogenic cell precursors. This study constitutes the first high-resolution, single-cell transcriptomic resource of somatic cell lineage commitment and differentiation during early ovarian development.

By combining single-cell RNA-sequencing of the XX and XY *Nr5a1*^+^ gonadal somatic cells from E10.5 to late developmental stages (E16.5 in XY, and P6 on XX), we provide the most comprehensive picture of cell lineage specification during gonadogenesis. We show that the supporting and the interstitial/stromal cell lineages both derive from a common *Nr5a1*^+^ early progenitor cell population, present at E10.5. We also demonstrate that the commitment of the supporting cells from the early progenitor cells is an active process and is disconnected from the initiation of sexual dimorphism, strictly speaking. However, the transient bipotential state of supporting cell precursors reveals a temporal asymmetry between XY and XX. In XY, this bipotential state corresponds to E11.5 pre-Sertoli cells, while in XX, it corresponds to E11.5 to E16.5 pre-granulosa cells (**Figure 6A**). The supporting cell precursors acquire their respective sex-specific genes from E12.5 onward in Sertoli cells, and from E13.5 onward in pre-granulosa cells. These results are consistent with the competence windows of supporting cells of both sexes to express *Sry* (Harikae et al., 2013). However, while the Sertoli cells completely down-regulate the bipotential supporting cell precursor genes, we found that the pre-granulosa cells maintain expression of stem cell-related genes until as late as E16.5, indicating that pre-granulosa cells remain in an early stage of their differentiation, and continue their differentiation from folliculogenesis onward (**Figure 6A**).

**Figure 6:**
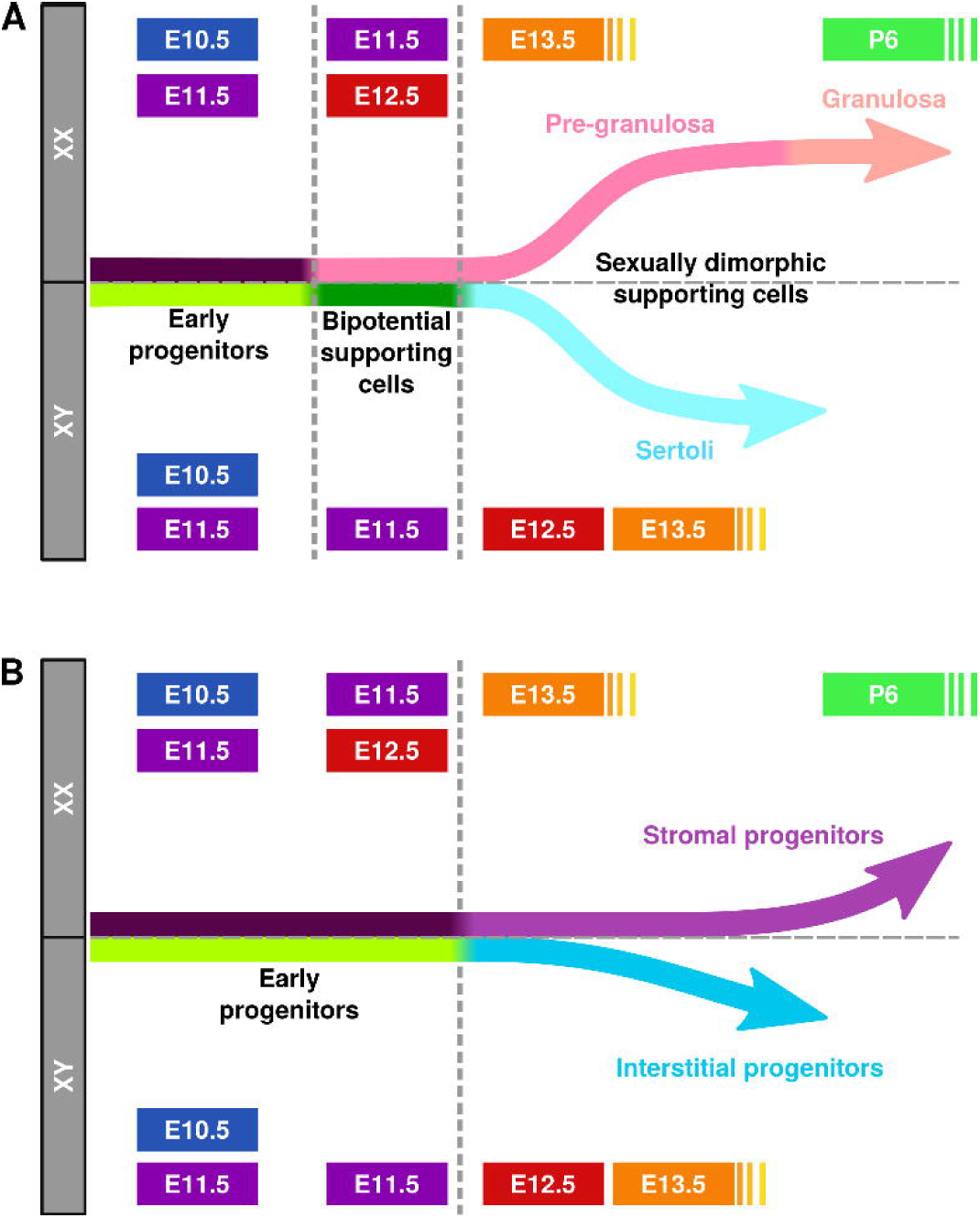
Model of somatic cell lineage commitment and sexual dimorphism acquisition in XX and XY during early gonad development. (A) Supporting cells differentiate from the early progenitors in two successive steps. The first step starts at around E11.5 with the commitment of the bipotential supporting cells that express a common genetic program, with the exception of *Sry* in XY cells. The supporting cells then undergo sexual specification, from E11.5-E12.5 subsequently to *Sry* expression in XY, and from E13.5 in XX. (B) The multipotent early progenitor cells undergo transcriptomic changes after the commitment of the supporting cells and become restricted to a steroidogenic cell fate. The interstitial and stromal progenitor cells do not display strong sexual dimorphisms, but differ in their timing.

Interstitial/stromal progenitor cells commit from early progenitors from around E12.5 in XY and around E13.5 in XX (**Figure 6B**). This temporal asymmetry is consistent with observations made in the supporting cell lineage. Surprisingly, interstitial/stromal progenitors do not display strong sexual dimorphism even late in embryonic development. Both XY and XX progenitor cells acquire a steroidogenic precursor fate by progressively expressing *Pdgfra, Arx* or *Ptch1*. The specification of the steroidogenic cell lineage is thus identical irrespective of the genetic sex of the individual, despite the differentiation of Sertoli and pre-granulosa cells. We hypothesize that supporting cells of both sexes control the specification of steroidogenic precursor cells in the same way, as they control the differentiation of theca and Leydig cells via the Hedgehog signaling pathway (Liu et al., 2015; Yao et al., 2002).

Our data provide a novel framework for further studies on the molecular and cellular programs of testis and ovary development. They also raise multiple questions, including the identity of the signals or factors that control the specification toward either the supporting or steroidogenic fate. Several parameters may influence this choice. First, it is possible that the fate of individual *Nr5a1*^+^ multipotent progenitors is affected by the local environment and interaction with neighboring cells. Second, it remains plausible that the multipotent progenitor population described here as a single population is in fact already heterogenous at the epigenetic level and composed of slightly different subpopulations. The epigenetic status and chromatin accessibility of these progenitor cells have not yet been investigated. Similar to embryonic stem cells (Atlasi and Stunnenberg, 2017), it has been hypothesized that the chromatin landscape in XX and XY progenitor cells of the gonad has an open configuration that confers on these multipotent cells a unique plasticity that enables differentiation into different lineages (Garcia-Moreno et al., 2018). Following cell fate commitment and sex-specific differentiation, the chromatin landscape becomes more restricted, canalizing the developmental program toward either the male or female fate and repressing the alternative pathway. We expect that the advent of new single-cell epigenomic sequencing methods, combined with single cell transcriptomic data, will be instrumental in advancing our understanding of the epigenetic regulation at play in each somatic cell lineage during mammalian sex determination.

## Experimental Procedures

### Mouse strains and isolation of purified *Nr5a1*-GFP positive cells

Animals were housed and cared for according to the ethical guidelines of the Direction Générale de la Santé of the Canton de Genève (experimentation GE-122-15). The experiment has been performed using heterozygous Tg(*Nr5a1*-GFP) transgenic male mice (Stallings, 2002). We performed the experiments in independent duplicates for each embryonic and postnatal stages. At each relevant gestation days, Tg(*Nr5a1*-GFP) gonads were collected and dissociated. Several animals from different litters were pooled together to obtain enough material for the experiment (**Table S1**). GFP^+^ cells were sorted by fluorescent-active cell sorting (**Figure S1**, **Supplementary experimental procedures**).

### Single-cell capture, cDNA libraries and sequencing

Cells were captured and processed using the C1 Autoprep System (96 well, small size chip), following the official C1 protocol. Sequencing libraries were prepared using the Illumina Nextera XT DNA Sample Preparation kit using the modified protocol described in the C1 documentation. Cells were sequenced at an average of 10 million reads (100bp paired-end reads) (**Supplementary experimental procedures**).

### Bioinformatics analysis

The computations were performed at the Vital-IT (http://www.vital-it.ch) Centre for high-performance computing of the SIB Swiss Institute of Bioinformatics. Data were analysed with R version 3.4.0. Cell clustering was performed on the highly variable genes with HCPC, differential expression analysis with Monocle2, diffusion maps were computed with Destiny, and lineage reconstruction was performed with Slingshot (for details, see **Supplementary experimental procedures**).

## Data and source code availability

XX single-cell RNA-seq data will be soon available on GEO. XY single cell RNA-seq data were taken from GEO (accession number GSE59698). Both XX and XY gene expression data are included in ReproGenomics Viewer (Darde et al., 2015).

## Author Contributions

Conceptualization, I.S., and S.N.; Investigation, I.S. and F.K.; Formal Analysis and Data Curation, I.S.; Writing – Original Draft, I.S., S.N., AG, and M.C.C.; Funding Acquisition, S.N., and E.T.D.; Resources, S.N., and E.T.D.; Supervision, S.N., and E.T.D.

## Acknowledgments

This work was supported by grants from the Swiss National Science Foundation (grants 31003A_173070 and 51PHI0-141994) and by the Département de l’Instruction Publique of the State of Geneva (to S.N.). We thank Luciana Romano and Deborah Penet for the sequencing. We thank also the members of the Nef and Dermitzakis laboratories for helpful discussion and critical reading of the manuscript, Cécile Gameiro from the flow cytometry facility for the cell sorting, and Didier Chollet, Brice Petit and Mylène Docquier from the iGE3 Genomics Platform for the single-cell capture.

---

**Supplementary Data 1: Differentially expressed genes between the 4 cell clusters, related to Figure 1H.**

**Supplementary Data 2: Biological Process GO terms of the differentially expressed genes between the cell clusters, related to Figure 1I.**

**Supplementary Data 3: List and classification of the genes showing a dynamic expression profile as the granulosa and the stromal progenitor cells differentiate, related to Figure 2C.**

**Supplementary Data 4: GO terms associated with the genes dynamically expressed during the granulosa and the stromal progenitor cell differentiation, related to Figure 3D.**

**Supplementary Data 5: List and classification of the genes showing a dynamic expression profile as the supporting cells from XX and XY gonads differentiate, and results of the differential expression analysis between the XX and the XY cells between the cell lineage branch points, related to Figure 4.**

**Supplementary Data 5: List and classification of the genes showing a dynamic expression profile as the progenitor cells from XX and XY gonads differentiate, and results of the differential expression analysis between the XX and the XY cells between the cell lineage branch points, related to Figure 5.**

